# Intravital imaging uncovers remodelling of humanised bone marrow-like niches

**DOI:** 10.64898/2026.03.24.713949

**Authors:** Colin D.H. Ratcliffe, Syed A Mian, Giovanni Giangreco, Alix Le Marois, Khadidja Habel, Erik Sahai, Dominique Bonnet

## Abstract

The bone marrow haematopoietic niche is composed of a diverse array of cell types and extracellular matrix components that together support healthy haematopoiesis. However, live imaging of the bone marrow microenvironment is hampered by tissue accessibility limitations. Using intravital imaging through a titanium imaging window, we investigated the dynamics of human haematopoietic cells and mesenchymal stromal cells within an ectopically implanted humanised scaffold in an immunodeficient murine host. These cell populations expand and differentiate over time, accompanied by progressive remodelling of the scaffold. We observe migration of murine endothelial cells into the scaffold, leading to the formation of a vascular network during the initial development of the humanised niche. Subsequently, the dense collagen matrix that makes up the implanted niche is altered and larger gaps form in regions populated by mesenchymal stroma cells. Collectively, our findings demonstrate dynamic remodelling of the extracellular milieu that supports haematopoietic cell development and establish a platform for longitudinal, in vivo investigation of these processes. Altogether, we describe a novel model that aligns with the 3R guiding principles and enables real-time assessment of bone marrow cell dynamics in vivo.

**Summary statement:** Ratcliffe and Mian et al. image *in vivo* dynamics of a bone marrow haematopoietic niche model.

## Introduction

Haematopoiesis is a tightly regulated and hierarchically organised process that sustains lifelong blood production and immune competence (Anjos-Afonso and Bonnet, 2023). In humans, it primarily takes place in the bone marrow, where haematopoietic stem cells (HSCs) reside within specialised microenvironments, or “niches,” composed mainly of mesenchymal stromal cells (MSCs), endothelial cells, osteoblasts, and other supportive cell types (Pinho and Frenette, 2019, Morrison and Scadden, 2014). These niches provide essential physical and biochemical cues that govern the quiescence, self-renewal, and lineage differentiation of HSCs. Understanding the intrinsic and extrinsic cellular as well as molecular mechanisms that regulate human haematopoiesis is crucial for developing therapies for haematological malignancies, improving transplantation outcomes, advancing regenerative medicine, and elucidating the interactions between human immune cells and solid tumour tissues.

Despite decades of progress, studying the complexity of human haematopoiesis in xenograft *in vivo* models remains challenging due to significant interspecies differences between humans and mice. To overcome these limitations, we and others have contributed to the development of ectopic humanised niches that are self-contained, human-derived haematopoietic microenvironments mimicking the human bone marrow tissue (Abarrategi et al., 2017, Mian et al., 2021, Reinisch et al., 2017, Ferrelli et al., 2025). In these humanised niches, human MSCs are integrated into biomaterial scaffolds to generate niches that can be implanted subcutaneously in immunodeficient mice. These scaffold-based niches reproduce critical features of the human bone marrow microenvironment, including cellular architecture, extracellular matrix composition and cytokine signalling. Crucially, they enable genetic manipulation of cellular components and end-point analyses *in vivo*. As such, humanised niche models have become powerful platforms for investigating HSCs, haematopoietic reconstitution, and therapeutic responses under both physiological and pathological conditions (Mian et al., 2020). However, we currently lack the ability to perform longitudinal monitoring of these ectopic humanised niches and their cellular components.

Imaging the bone marrow niche *in vivo* has become an increasingly powerful approach, as studies demonstrate that niche architecture and dynamic cell–cell interactions critically influence stem cell fate and behaviour (Calvi and Link, 2015, Pinho and Frenette, 2019, Chacon-Martinez et al., 2018). Intravital two-photon microscopy enables the study of cellular dynamics within living tissues. With high spatial and temporal resolution, it is ideally suited for visualising the behaviour of cells within their native environments. We have developed an imaging protocol combining fluorescent labelling of human cells and a custom-designed mouse imaging stabilisation apparatus (MISA) that enables intravital microscopy through an implanted titanium window. This approach enables visualisation of the dynamics of human haematopoietic cells as well as the niche microenvironment. It also gives us the ability to compare the localisation of cellular components with the altered niche stromal components.

Together, the integration of ectopic humanised niche model with advanced intravital two-photon imaging provides an unprecedented opportunity to investigate human haematopoiesis *in vivo* at a high resolution. These complementary methodologies may be used to enhance our understanding of human haematopoietic stem cell dynamics within a physiologically relevant and experimentally accessible context.

## Results

### Establishing humanised niches with titanium-based imaging window in immunodeficient mice

We have previously established an ectopic *in vivo* environment that provides a physiologically relevant niche capable of supporting human cell attachment, organisation, and the development of bone-marrow–like tissue (Abarrategi et al., 2017, Mian et al., 2021). To explore the dynamics of this environment, we incorporated additional steps in which human haematopoietic stem and progenitor cells (HSPCs) and mesenchymal stromal cells (MSCs) were lentivirally labelled with a fluorescent reporter and pre-seeded into collagen-based porous scaffolds—collectively termed the humanised ectopic niche scaffold model. This protocol supports the formation of ectopic niches containing labelled human cells onto which a titanium imaging window could be surgically implanted (**Fig 1A**).

**Fig. 1.**
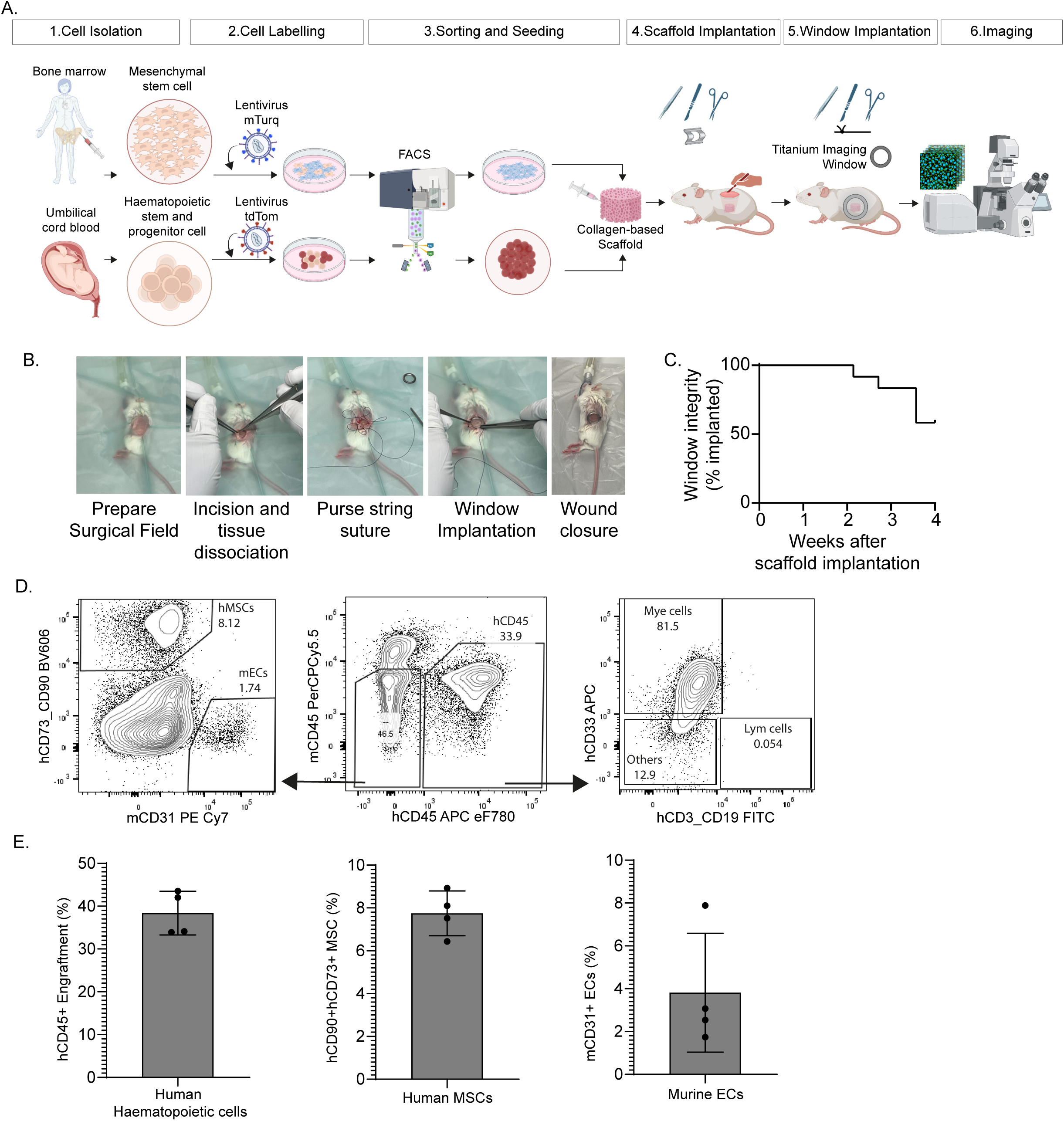
Establishment of a bone marrow-like niche model amenable to intravital imaging (A) **(A)** Schematic of experimental steps and timeline required for implantation and imaging of humanised scaffold. **(B)** Representative photographs of the surgical steps required for implantation of imaging window. **(C)** Probability curve showing imaging window retention following surgical implantation. Data represent a total of 12 mice included in the analysis. **(D)** Representative flow cytometry dot plots showing human haematopoietic cells, MSCs and mouse endothelial cells at four weeks after the window implantation (or 5 weeks after scaffold implantation). **(E)** Bar graphs quantifying the engraftment of human haematopoietic cells, MSCs and mouse endothelial cells at four weeks post-window implantation (or five weeks after scaffold implantation).

We first sought to compare two implantation strategies. A co-implantation approach of scaffolds pre-seeded with HSPCs and MSCs simultaneously implanted with the imaging window was evaluated against a sequential approach in which scaffolds were established before subsequent window placement. Three weeks after window implantation, mice were culled and the scaffolds retrieved for analysis. Although the scaffolds were macroscopically comparable, flow cytometry revealed markedly higher human cell engraftment in the serial implantation group (human CD45⁺ cells, 39.1% vs 2.89%) (**Supp. Fig. 1**). Based on this observation, we used the serial implantation approach for all the subsequent experiments.

Next, we assessed the stability of the window over a timeframe that would be consistent with the development of the humanised niche. First, Human HSPCs and MSCs were transduced with a leGO-IT reporter system expressing tdTomato and mTurquoise respectively (Weber et al., 2008). Fluorescently labelled cells were flow-sorted, and pre-seeded into collagen-based scaffolds. A single scaffold was implanted into each NSG-SGM3 mouse and one week later, a titanium-based imaging window was implanted (n=12) (**Fig. 1B, Supplementary video 1**). Mice were then monitored longitudinally to assess window retention and niche formation. Most imaging windows remained securely attached for three weeks (>80%), and 58% (7/12) were retained at four weeks (**Fig. 1C**). Flow cytometry analysis of retrieved scaffolds after the window implantation confirmed robust engraftment of both human haematopoietic cells and MSCs. Importantly, the presence of the imaging windows did not impede the engraftment of both human cell types. In addition, murine endothelial cells were detected within fully developed scaffolds, indicating successful vascular integration (**Fig. 1D-1E**). These results are consistent with our previous reports (Abarrategi et al., 2017, Mian et al., 2021) and demonstrate that window implantation maintains the establishment, vascularisation, and structural integrity of the humanised niches.

### Visualization of humanised niches using two-photon microscopy

In vivo imaging of the bone marrow niche has emerged as a powerful strategy to interrogate how microenvironmental architecture and dynamic cell–cell interactions regulate stem cell fate and behaviour (Calvi and Link, 2015, Pinho and Frenette, 2019, Chacon-Martinez et al., 2018). Intravital microscopy uniquely permits direct, real-time visualisation of these processes within their native physiological context, preserving spatial organisation and vascular integrity. To enable intravital imaging of the humanised niche, we custom-designed a mouse imaging stabilisation apparatus (MISA) (**Fig. 2A**) to securely position live mice on an upright microscope and minimize motion caused by breathing during isoflurane-induced sedation. The MISA comprises three main components, which attach to the microscope’s imaging platform and are secured together with six bolts. The device also features a liquid-holding chamber positioned above the mouse, providing the necessary interface between the 1.05 N.A. water objective and the implanted window, thereby ensuring optimal imaging conditions (**Fig. 2A-2B**).

**Figure 2:**
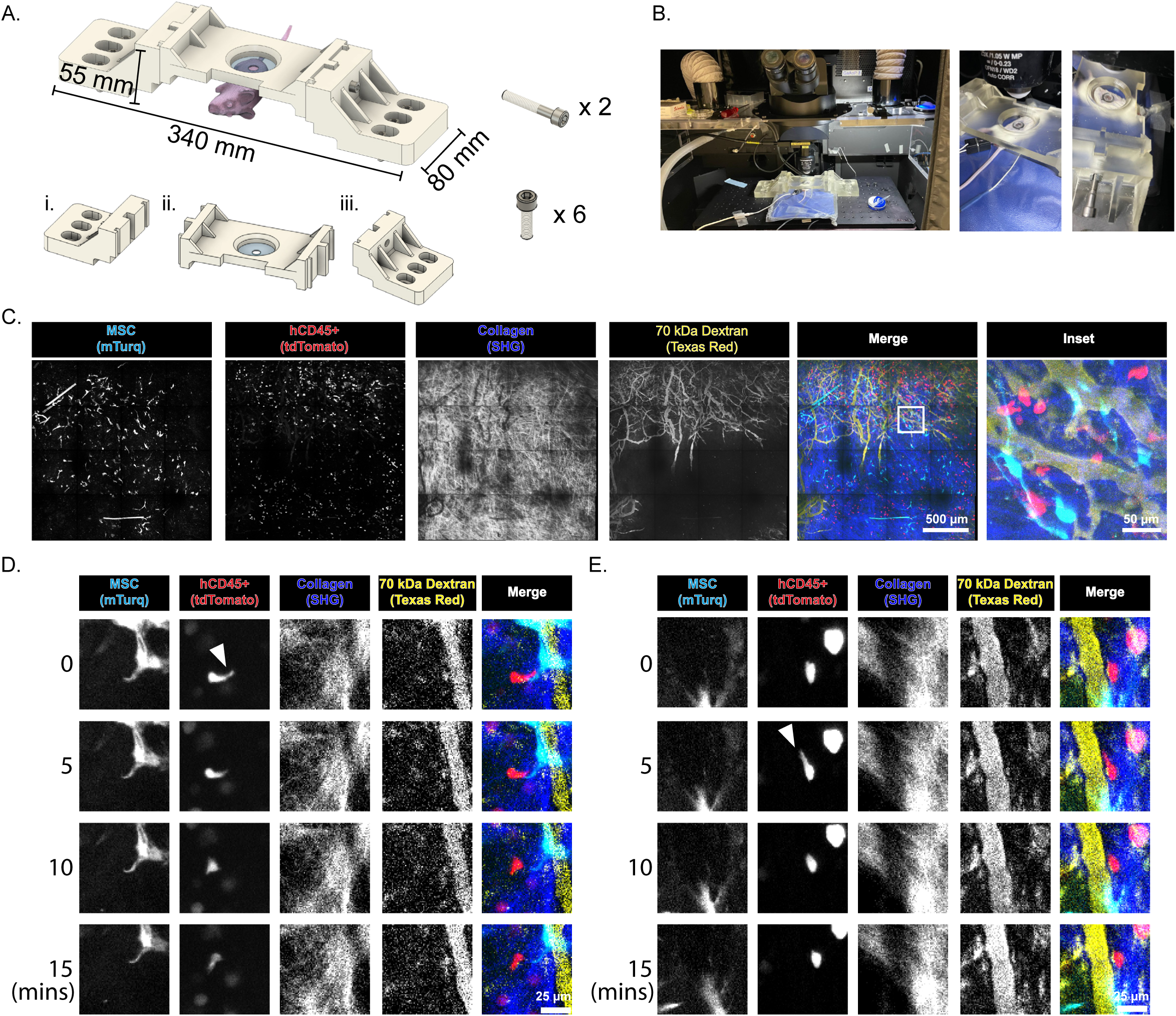
Imaging windows enable investigation of the humanised niche in mice. **(A)** Schematic of mouse imaging stabilising apparatus. The size of the device is 340mm in length, 80mm in width and 55mm in height. **(B)** Photographs of the microscope setup with the MISA and a mouse prepared for intravital imaging. The assembled MISA securely holds the mouse in place, ready for intravital imaging **(C)** Representative micrographs of humanised scaffold with human MSCs (cyan), haematopoietic cells (red) and collagen (blue). Mice were injected with Texas red labelled 70kDa dextran (yellow) immediately prior to imaging. Right-hand panels show merged images of all cell types within the humanised niche. **(D-E)** Selected micrographs from time lapse imaging of a single field of view within the humanised niche. Imaging was performed over the course of 15 minutes. Right-hand panels show merged images of all cell types within the humanised niche.

Using an Olympus FV3000 system connected to two laser lines (one variable line set to 820 nm and a fixed line at 1045 nm), we could readily detect mTurq labelled MSCs, tdTomato HSCs, collagen structure and the murine vascular network (**Supplemental Fig. 2)**. Beam splitters allowed us to image the collagen matrix using second harmonic generation and excite mTurq with the 820 nm laser. Simultaneously, the 1045 nm laser line was used to excite tdTomato along with Texas Red conjugated to 70 kDa dextran that had been injected into the tail vein prior to imaging (**Fig. 2C & Supplemental Video 2**). Altogether, our protocol enables *in vivo* imaging of 4 components of the humanised scaffold across a 3.9 mm^3^ field of view.

Cell motility is a defining feature of haematopoietic biology and has been widely described in mouse models and patient-derived xenograft models, particularly within the native bone marrow where haematopoietic cells exhibit active migration, perivascular patrolling, and stochastic displacement (Lo Celso et al., 2009, Sipkins et al., 2005, Foster et al., 2015). To assess whether such behaviour is retained in our humanised ectopic niches, we imaged both cell types and extracellular components of the niche over a 15-minute interval. Within the imaged regions, tdTomato+ cells displayed motile behaviour, including reorientation through the scaffold matrix and movement away from perivascular zones **(Figure 2D & Supplemental Video 3)**. We observed haematopoietic cells positioned along vessel structures (**Fig 2E &, Supplemental Video 4**). This dynamic behaviour recapitulates the interactions and motility patterns described in native murine bone marrow environments (Foster et al 2015).

### Tracking humanised niche maturation via longitudinal imaging

Tissue niches are inherently dynamic structures that undergo continuous remodelling as their cellular components interact with and reshape their surrounding microenvironment (Winkler et al., 2020, Wang and Wagers, 2011). Likewise, the ectopically implanted humanised niche scaffold is a dynamic structure that undergoes maturation as murine components integrate with it and human components expand and remodel the tissue. We sought to visualise this gradual process through longitudinal intravital imaging. As the imaging windows remained stably positioned over the humanised scaffolds for up to four weeks (**Fig. 1C**), we selected a three-week imaging period to ensure reliable, repeated high-resolution visualisation of the regions within these niches.

An added benefit of the staggered-window implantation strategy is that it enables us to perform longitudinal monitoring of both early and later stages of niche development. We synchronised initiation of niche development by implanting pre-seeded scaffolds into mice (n = 12), and imaging windows were surgically placed in subsets of mice (n = 4) at 2-, 5-, and 8-weeks following scaffold implantation. Imaging mice between weeks 3 and 5 post implantation revealed early phases of niche establishment. Across this interval, we observed increases in both haematopoietic cells and MSCs within the visualised regions of the humanised scaffolds. Notably, the most pronounced expansion occurred between weeks 3 and 4, during which there was a rise in the number of haematopoietic cells and MSCs (**Fig. 3A-3B**).

**Figure 3:**
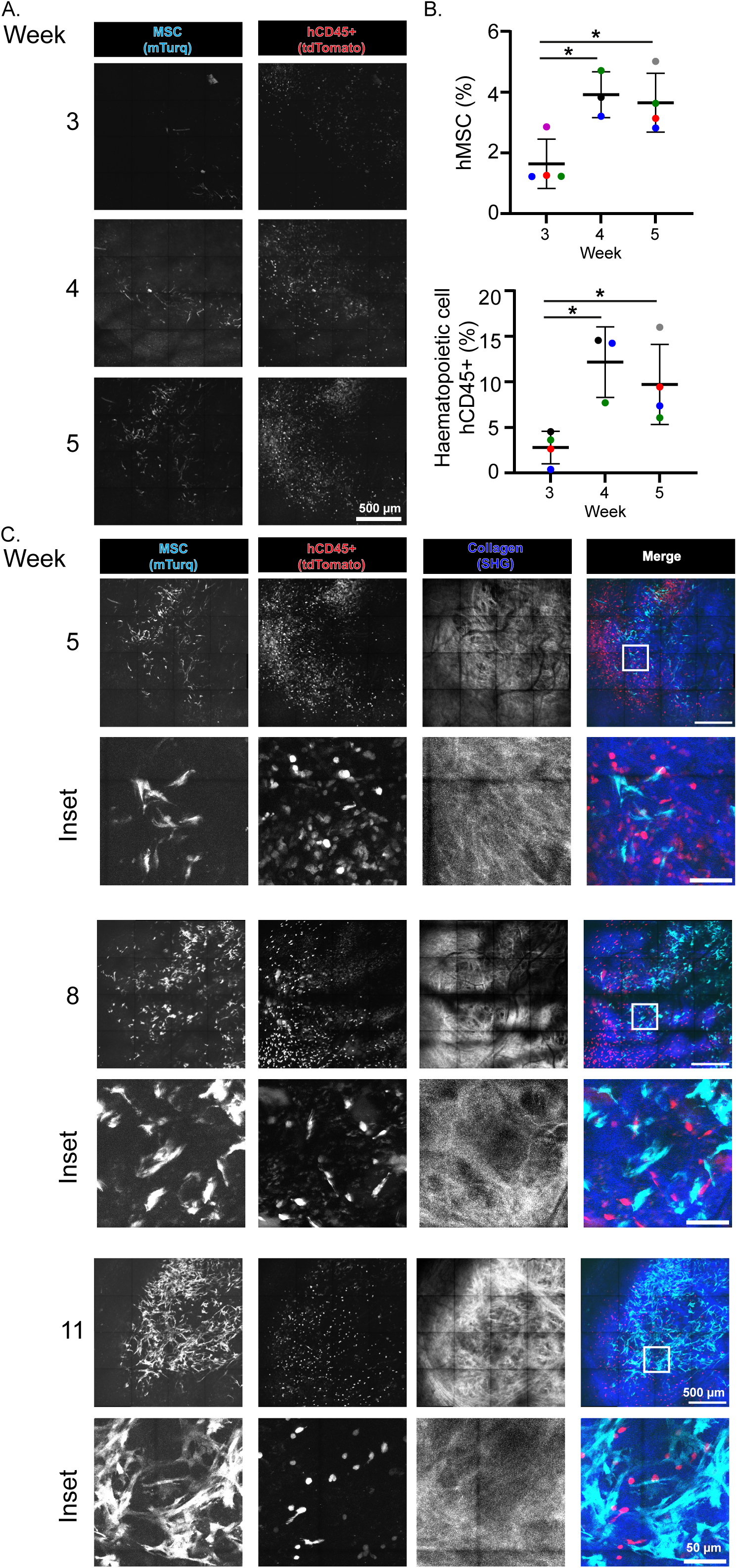
Visualising MSC and haematopoietic cell expansion in the humanised niche. **(A)** Representative micrographs of haematopoietic cells and MSCs in humanised scaffolds from the same mouse. Intravital imaging was performed at weeks 3, 4, and 5 post implantations of the imaging windows. Right-hand panels show merged images of all cell types within the humanised niche. **(B)** Intraviral imaging-based quantification of human haematopoietic cells and MSCs within the imaged region of the humanised scaffold at different time points following the window implantation. Datapoint colour correspond to mouse identity **(C)** Representative micrographs of haematopoietic cells, MSCs and collagen in humanised scaffolds at weeks 5, 8 and 11 post implantations. For each image, inset is indicated by the white box. Right-hand panels show merged images of all cell types within the humanised niche. * p <0.05

At later stages of niche development (weeks 8 and 11), the scaffolds were characterised by haematopoietic cells and MSCs interspersed throughout (**Fig. 3C**). This is consistent with reports that early niche maturation is driven by simultaneous stromal proliferation and haematopoietic cell engraftment, leading to the establishment of more densely populated and functionally organised microenvironments (Sanchez-Lanzas et al., 2024, Mesnieres et al., 2021, Coskun et al., 2014, Cain et al., 2025).

### Temporal dynamics of collagen matrix

Interestingly, we observed different collagen structures at these later time points compared to earlier ones. Collagen structures provide essential physical cues for stem cells throughout the body including in the bone haematopoietic niche (Pinho and Frenette, 2019, Lee-Thedieck et al., 2022). We generated another cohort of mice and imaged these 5-, 6- and 7-weeks post scaffold implantation and implemented our framework for fibre analysis, termed TWOMBLI (Wershof et al., 2021), to quantify the changes in collagen organisation (**Fig. 4A**). As time proceeded, we performed SHG of collagen scaffolds and analysed metrics. At week 5, scaffolds appeared dense, with smaller gaps between fibres. After 7 weeks post implantation of the collagen scaffold, we observed no changes in the frequency of branch points along the collagen scaffold (**Fig. 4B**). Trends towards decreased fibre alignment and fraction of high-density matrix were not significant (**Fig. 4C-D**). However, large gaps were readily apparent in the matrix and these made up a significantly larger proportion of total gaps (**Fig. 4E**). These observations support a model whereby the collagen scaffold, not only provides structural signals for haematopoietic niche development but is actively remodelled.

**Figure 4:**
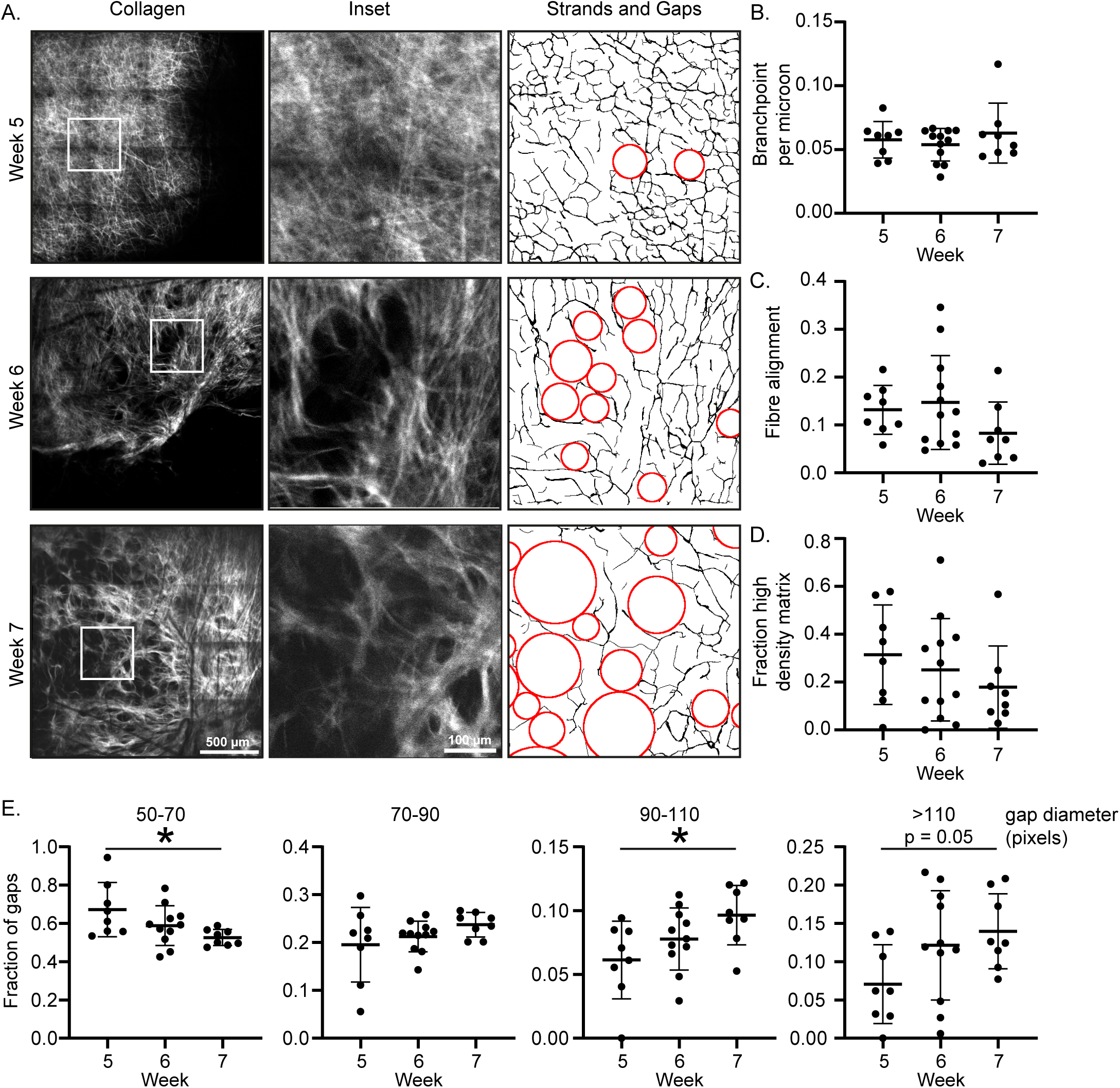
Scaffold collagen is remodelled over time. **(A)** Representative micrographs and insets of collagen in humanised scaffolds. Intravital imaging was performed at weeks 5-, 6- and 7- post window implantation. Right-hand side of the panel is an image output of TWOMBLI gap analysis showing collagen strands (black lines) and gaps (red circles) (> 50 pixels). **(B, C, D, E)** Intravital image-based analysis of the data derived from TWOMBLI including (B) Branchpoints per micron, **(C)** Fibre alignment and **(D)** fraction of high-density matrix. **(E)** Frequency analysis of gap sizes in collagen matrix of humanised scaffolds. * p <0.05

While gaps were present at week 7, they were not uniform across the scaffold, and we observed regions that had been remodelled while others appeared dense. We hypothesised that the human cellular components of the niche are involved in this remodelling. Using our simultaneous multi-channel imaging, we computed the colocalization coefficient of MSCs on gaps for each image (termed ρ_MG_), and found that the degree of colocalization between MSCs and gaps increased between weeks 5 and 7 (**Fig. 5A**).

**Figure 5:**
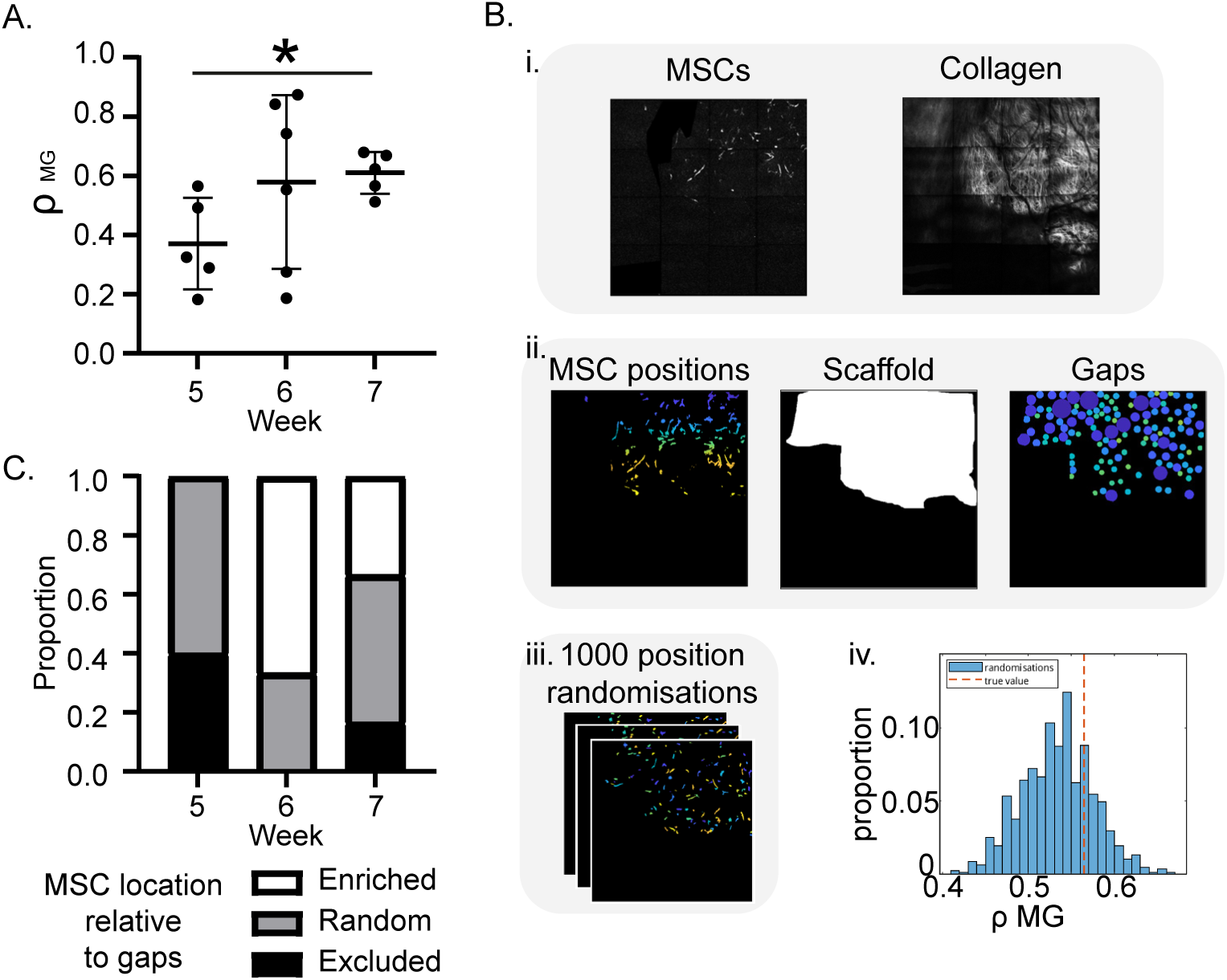
Gaps in collagen scaffolds are enriched for MSCs. **(A)** Quantification of the colocalization of MSCs with gaps at weeks 5-, 6- and 7- post window implantation. **(B)** Diagram of the steps taken to quantify the enrichment of MSCs on gaps relative to random. i. Images of MSCs and collagen ii. Masks of MSCs where colours correspond to MSC identity, scaffold mask and gap mask where colours correspond to gap size. iii. Representative images of MSCs randomised within the scaffold space. iv. Representative histogram showing the proportions of images with varying the ρ_MG_ values and the true value (red line). **(C)** Bar chart for each time point showing the proportion of images where MSCs were either enriched, randomly distributed or excluded from gaps. * p <0.05

To confirm that the enrichment of MSCs on gaps over time was not due to the increase in gap size, for each image, we generated a series of 1000 synthetic masks where the positions of the original MSC cells were randomly distributed within the scaffold area (**Fig. 5B**). We then computed the colocalization coefficient for each of the 1000 randomised images (ρ’_MG,i_ with i=1->1000). Images for which ρ_MG_ fell within the distribution of ρ’_MG,i_ values were categorised as showing a proportion of MSCs on gaps consistent with random distribution. Images where ρ_MG_ was higher or lower than 95% of the ρ’_MG,I_ values were categorised as showing significant enrichment or exclusion of MSCs on gaps, respectively.

Using this approach, at week 5, we observed random distributions of MSCs on gaps or exclusion from gaps, with no images showing enrichment. In weeks 6 and 7, fewer images showed exclusion, while several images showed enrichment of MSCs on gaps (42% at week 6 and 28.5% at week 7) (**Fig. 5C**). This suggested a progressive overall concentration of MSCs in remodelled areas of the scaffold over time. The lower proportion of images with enrichment at week 7 compared to week 6 could be due to a large increase in the number and size of gaps observed in the scaffold at week 7, meaning randomly distributed cells are highly likely to localise on gaps, and higher thresholds were required for co-localisation to be classified as enrichment.

These data uncover a connection between scaffold remodelling and mesenchymal stroma cells in a haematopoietic niche. Altogether they support a model whereby ectopic humanised niches in immunodeficient mice provide a powerful system for investigating human bone marrow biology *in vivo* (Reinisch et al., 2016, Abarrategi et al., 2017, Ferrelli et al., 2025, Mian et al., 2021, Grigoryan et al., 2022).

## Discussion

Here, we introduce a longitudinal imaging platform for ectopically implanted humanised niches, which enables direct studying of dynamic cellular behaviours and interactions within these niches—processes that have previously remained difficult to investigate.

By integrating fluorescently labelled human haematopoietic cells within a collagen-based scaffold and overlaying a titanium imaging window, we achieved repeated high-resolution imaging without compromising tissue development or cellular viability. This configuration supported robust engraftment of human cells *in vivo* while maintaining continuous optical access. Importantly, flow cytometric analyses of xenografted cells confirmed that humanised niche development occurred in the presence of the imaging window, validating the platform’s suitability for functional and mechanistic studies. Imaging windows remained stably positioned for up to four weeks, and the custom-designed MISA effectively minimised motion artefacts when imaging. Our approach enabled us to study the dynamic remodelling of humanised niches, driven by coordinated interactions among haematopoietic cells, stromal cells, the vasculature, and the surrounding ECM.

We observed haematopoietic cell movement and spatial organisation within the humanised niches. Traditional approaches rely on static, endpoint analyses that have limited our ability to capture temporal dynamics. In concordance with previous reports, murine endothelial cells infiltrated the humanised scaffold during early development and established a functional vascular network (Bessy et al., 2021), reflecting host-driven vascularisation process. Human MSCs expanded within these developing vascular structures, while haematopoietic cells could be observed dynamically interacting with MSCs and vascular structures. Therefore, the imaging protocol for ectopically implanted humanised niches permits observation of cellular behaviour over biologically relevant time scales, providing insight into processes that underpin functional bone marrow haematopoiesis.

We then used our system to track niche maturation over several weeks. Temporal analyses of the imaged areas of the humanised niche revealed progressive expansion of human haematopoietic cells and MSCs accompanied by pronounced collagen remodelling during niche development, reflecting a transition from initial engraftment toward an established mature, functional tissue. These dynamics parallel processes observed in native bone marrow (Pinho and Frenette, 2019), where coordinated stromal remodelling, vascular integration, and ECM maturation underpin the establishment of sustained haematopoiesis.

The physiological bone marrow environment in which MSCs reside is characterised by a complex ECM rich in multiple collagen types, fibronectin, and laminin, even within the densely cellular perivascular niche. In native human bone marrow, type I collagen predominates at bone surfaces providing structural rigidity, whereas type III collagen forms a reticular network within the marrow stroma, supporting haematopoietic cells. SHG imaging revealed collagen remodelling, increasing gap sizes and enrichment of MSCs in these spaces between 5- and 7-weeks post-scaffold implantation. Previous studies indicate that MSC-deposited, collagen-rich ECM provides not only structural support but also essential biochemical and biophysical cues that regulate cell adhesion, migration, and fate decisions (Watt and Huck, 2013, Guilak et al., 2009). In particular, interactions between haematopoietic cells and collagen matrices, mediated through integrin-dependent signalling pathways, have been implicated in maintaining stem cell localisation, quiescence, and lineage specification (Pinho and Frenette, 2019, Lee-Thedieck et al., 2022, Chen et al., 2013, Donnelly et al., 2024). Our observations support this paradigm, where ECM remodelling within the humanised niche, likely involving both type I– and type III–like collagen structures, could be actively contributing to mechanotransductive signalling and spatial organisation.

Together, our findings establish the imaging platform for ectopically implanted humanised niches as a translationally relevant system for interrogating human haematopoietic niche biology *in vivo*. By enabling continuous, high-resolution visualisation of humanised bone marrow–like tissues, we provide methodology to assess the influence of genetic perturbations, pharmacological agents, or engineered biomaterials on stem cell behaviour and niche function over time. Additionally, our approach eliminates the need for multiple experimental cohorts and reduces inter-animal variability thereby, aligning with the principles of the 3Rs (Replacement, Reduction and Refinement). Moreover, we envision this platform can be readily adapted for preclinical testing of niche-targeted therapies, optimisation of cell-based manufacturing protocols, and assessment of human stem cell engraftment dynamics in response to microenvironmental cues. Further studies incorporating direct comparisons with human bone marrow trephine biopsies will be essential to evaluate how closely the spatial organisation, collagen distribution, and niche compartmentalisation of our humanised constructs reflect physiological human bone marrow. As such, the humanised niche imaging system bridges a critical gap between reductionists in vitro models and endpoint-driven in vivo assays, providing a versatile and scalable tool for translating fundamental insights in haematopoiesis and niche biology into clinically actionable strategies.

## Materials and Methods

### Bone marrow samples and umbilical cord blood cells

Bone marrow mononuclear cells (MNCs) were obtained from Stemcell Technologies and stored at the Francis Crick Institute. Umbilical cord blood cells were purchased from Anthony Nolan. All donors provided written informed consent in accordance with local tissue bank regulations, and all research procedures were conducted in compliance with the Declaration of Helsinki.

### MSC isolation and expansion

Human bone marrow MNCs were used for MSC isolation and expansion. Briefly, CD34^+ve^ selection was performed using Easysep human CD34 positive selection kit (StemCell Technologies,) and Easysep magnet (StemCell Technologies) according to the manufacturer’s instructions. CD34^-ve^ cells were seeded at a density of 1×10^6^/cm^2^ using MSC culture media [MEM Alpha Medium (1X) + GlutaMAX-1 (Gibco, Cat 32571–029), 10% human MSC-FBS (Gibco) and 1% penicillin/streptomycin (Sigma-Aldrich]. After 24 hours of incubation, the MSC culture media was replaced to remove non-adherent cells, and adherent MSCs were washed with PBS before adding fresh MSC culture media. The culture medium was subsequently refreshed once per week. Expanded MSCs were cryopreserved as viable cells at passage 1.

### CD34^+^ isolation from the umbilical cord blood

Mononuclear cells (MNCs) from UCB samples were isolated by density centrifugation using Ficoll-Paque TM PLUS (GE Healthcare Life Sciences, Buckinghamshire). Red blood cells were lysed by incubating the cells with ammonium chloride solution at 4°C for 10 minutes. Subsequently, CD34⁺ haematopoietic stem and progenitor (HSPCs) were enriched using the EasySep™ Human CD34 Positive Selection Kit II (STEMCELL Technologies), following the manufacturer’s instructions. Individual UBC samples from individual donors were pooled together according to donor sex.

### Lentivirus production, purification, and titration

The LeGO-tdTom plasmid (Addgene plasmid #27342) (Weber, Bartsch et al. 2008) was purchased from Addgene. The LeGO-mTurquoise2 plasmid was kindly provided by the laboratory of Boris Fehse. Briefly, the mTurquoise2 cDNA was derived from the plasmid pPalmitoyl-mTurquoise2 (Addgene plasmid #36209) and used to replace the eGFP cDNA in the LeGO-G2 (Addgene plasmid Plasmid #25917) backbone. The mTurquoise2 sequence was PCR-amplified using the primers (p219-Fw: ATAGGATCCCGCCACCATGGTGAGCAAGGG p220-Rv: ATAGAATTCTTACTTGTACAGCTCGTCCATGCC) and subsequently cloned into the LeGO vector using the BamHI and BsrGI restriction sites. Lentiviral particles were produced by transient CaCl₂-mediated transfection of human embryonic kidney 293 cells (HEK-293) with LeGO plasmids together with the packaging plasmid psPAX2 (psPAX2 was a gift from Didier Trono, Addgene plasmid Plasmid #12260) and the envelope plasmid pMD2.G (pMD2.G was a gift from Didier Trono, Addgene plasmid # 12259). HEK-293 cells were maintained in Dulbecco’s modified Eagle medium (DMEM) supplemented with 10% fetal bovine serum (FBS; Sigma-Aldrich) and 1% penicillin–streptomycin during the first 24 hours post-transfection. Twenty-four hours later, the culture medium was replaced with serum-free Opti-MEM (Thermo Fisher Scientific). Viral supernatants were harvested after an additional 24 hours, filtered, and concentrated by ultracentrifugation (28,000 × g for 2 hours). Viral titers were determined, and transduction efficiency was assessed by flow cytometry based on the proportion of infected cells.

### Lentivirus-based transduction of human cells with LeGO plasmids

Human UCB CD34⁺ HSPCs were pre-stimulated in StemSpan medium (STEMCELL Technologies) supplemented with cytokines including stem cell factor (SCF, 150 ng/mL), interleukin-6 (IL-6, 10 ng/mL), Flt-3 ligand (150 ng/mL), granulocyte colony-stimulating factor (G-CSF, 25 ng/mL), and thrombopoietin (TPO, 20 ng/mL) (PeproTech), as well as 1% HEPES (Sigma-Aldrich), for 4–6 hours. Following stimulation, tdTomato lentiviral particles were added at a multiplicity of infection (MOI) of 30, and cells were incubated overnight at 37°C. HSPCs were then washed and resuspended in expansion StemSpan medium that was supplemented with cytokines [SCF (150 ng/ml), IL-6 (10 ng/ml), Flt-3 (150 ng/ml), G-CSF (25 ng/ml), and TPO (20 ng/ml)] and 1% HEPES. Cells were expanded for 4 days prior to fluorescence-activated cell sorting.

Human bone marrow–derived MSCs were counted and seeded into 6-well plates, then transduced with mTurquoise lentiviral particles at an MOI of 10. Twenty-four hours post-transduction, cells were washed with PBS and cultured in fresh MSC growth medium. MSCs were expanded for 10 days before being subjected to fluorescence-activated cell sorting.

### Flow cytometry and cell sorting

Transduced MSC and UCB CD34^+^ enriched cells were stained with DAPI (4,6, diamidino-2-phenylindole, 1:2000 dilution from a 200 g/ml stock) to distinguish viable from non-viable cells. Cell sorting was performed on a FACSAria™ SORP (BD Biosciences, Oxford, UK). After exclusion of debris, doublets, and dead cells, CD34⁺tdTomato⁺ HSPCs and mTurquoise⁺ MSCs were isolated.

Post-sorting, MSCs were further expanded to passage 3 before use in scaffold seeding, while sorted HSPCs were used directly for scaffold seeding.

### Cell seeding into the Collagen-based scaffolds

Humanised scaffolds were prepared as previously described (**Fig 1A**) (Mian et al., 2021, Mian and Bonnet, 2021). Briefly, mTurquoise⁺ MSCs (100,000 cells) were seeded into the scaffolds and cultured for 48 hours. Subsequently, tdTomato⁺ HSPCs (30,000-40,000 cells) were introduced into the scaffolds. The scaffolds were then incubated for 24 hours in H5100 medium (STEMCELL Technologies) to be ready for implantation into mice the following day.

### Surgical implantation of the humanised scaffold

Surgical implantation of the pre-seeded scaffolds was performed in accordance with aseptic techniques as outlined by local veterinary surgical guidelines (as described previously (Mian et al., 2021, Mian and Bonnet, 2021)). NSG-SGM3 mice were used for all procedures. The surgical area (left or right flank) on the mouse was shaved 24 hours prior to surgery, and mice were provided with carprofen (0.1 mg/mL) in drinking water as pre-emptive analgesia. On the day of surgery, mice were anesthetized in an induction chamber containing 5% isoflurane in 1.5 L/min oxygen. Additional analgesia [buprenorphine (0.1 mg/kg) and meloxicam (10 mg/kg)] was administered subcutaneously.

The surgical site was sterilized using a 10% chlorhexidine solution. A 0.5cm incision was made in the anterior–posterior direction. A subcutaneous pocket was created using blunt-ended scissors (or forceps), and the pre-seeded scaffold(s) were inserted deep into the pocket. The incision was closed with surgical staples. Post-operative analgesia (carprofen, 0.1 mg/mL in drinking water) was maintained for 48 hours following surgery. Surgical staples were removed 7–10 days post-surgery.

### Imaging window implantation

To make implantable imaging windows, cover glass (round, 12 mm, No. 1.5, Paul Marienfeld ref 0117520) was super glued to implantable titanium ring and, following drying, these were sterilised by ethylene oxide gas. Surgical implantation was performed in accordance with aseptic techniques as outlined by local veterinary surgical guidelines. The surgical area was shaved within 24 hours prior to surgery and mice were provided with carprofen (0.1 mg/mL) in drinking water. On the day of surgery, mice were anaesthetised in an induction chamber containing 5% isoflurane in 1.5 L/min oxygen. Analgesia [buprenorphine (0.1 mg/kg) and meloxicam (10 mg /kg)] was administered subcutaneously.

The surgical site was sterilised using a 10% chlorhexidine solution and a 2-2.5cm incision was made in the anterior-posterior direction immediately dorsal of the scaffold with the scaffold at the midpoint of the incision. The imaging window was placed and secured using Mersilk 4-0 (1.5 Ph. Eur) C-3 13 mm 3/8c reverse cutting needles. Three triple knots in alternating directions were used to secure the window. Post-operative analgesia (carprofen, 0.1 mg/mL in drinking water) was maintained for 2 days following surgery.

### Design and printing of MISA

The device was designed using Alibre Design software (version 21). The apparatus parts and o-rings were printed with Preform Form3B printer with 0.1 mm layer height. Clear V4 resin was used for the platform and Elastic V1 resin was used for the o-rings. The apparatus was held together by six M6 x 16mm full thread socket cap head screws (ACCU SSCF-M6-16-A2), two M6 x 35mm socket cap head screws (ACCU SSC-M6-35-A2) and two M6 self-tapping inserts (ACCU HSTI-M6-A2). Autodesk Fusion 360 (version 2605.1.52) was used to create Figure 2A.

### Two-photon live imaging of the humanised niches

Using the MISA, pieces i and iii were fastened to the stage using 3 screws each. A far infrared heating pad along with a temperature probe was covered in plastic wrap and placed between these two pieces. Mice were anaesthetised using isofluorane and then placed over the heating pad and a small amount of Vaseline was added to the perimeter of the titanium ring. A pulse oximeter paw sensor was then placed on a lower hind paw to monitor the mouse using a PhysioSuite (Kent Scientific Corporation Item ID PS-03). A small amount of Vaseline was added to the perimeter of the hole of piece ii and then it was placed over the mouse. Piece ii was lowered to create a seal with the imaging window as well as immobilise the mouse. It was then fastened in place with a screw on each side. A drop of water was added over the window.

Intravital images were acquired on an Olympus FV3000 MP-RS microscope connected to a Spectra Physics Insight X3 (with variable line (690nm-1300nm) and fixed line at 1045nm. Emission light was collected with two PMT and two GaAsP non-descanned detectors. The objective used was the XLPLN25XWMP2 1.05 NA (water). A SDS595LP beam splitter was used to send reds onto the PMT and blue/greens onto the GaAsP detectors. Reds were passed through the Chroma T595lpxr beamsplitter and Semrock FF01-624/40 or Semrock FF02-575/25 emission filters. Blue/greens were passed through a Chroma T425lpxr beamsplitter and Semrock FF01-406/15 or Semrock FF02-472/30 emission filter. Image sizes were 1976 x 1976 pixels at a resolution of 0.994 micron/pixel. 40 images were captured per z-stack at two-micron intervals. Most images in this study were generated from 4 x 4 tile scans. Time lapse images were taken every 30 seconds over the course of 15 minutes. Resonant scanning was used on roundtrip scanning direction, pixel dwell time was 0.067 microseconds, lines were acquired sequentially, and integration type was set to frame. Integration count was set to 8. The microscope was controlled using Olympus FLUOVIEW FV315-SW software.

### Image processing and analysis

Tiled images were stitched using Olympus FLUOVIEW FV315-SW and later processed using FIJI. Out of 40 z-positions captured, 20 consecutive frames from each mouse and time point were selected for further analysis. Where indicated either sum intensity or maximum intensity projections were generated. For analysis, regions of interest were generated by manually outlining scaffolds based on second harmonic imaging. Collagen fibre and gap analysis was performed using TWOMBLI 2.0 (Wershof et al., 2021) (GitHub).

To quantify the distribution of MSCs within the scaffold area, we used images generated of the gap analysis and MSCs were segmented using cellpose with the following parameters: cyto3 model, input diameter = 30, cell probability threshold = -2 and flow threshold : 0. Intensity- and area-based filtering was then applied to remove small and dim non-cellular objects. A cell was said to localise on a gap if at least 10% of its pixels overlapped with a gap. For each image, the proportion of MSC cells on gaps was computed based on these criteria, yielding the co-localisation coefficient ρ_MG_.

Since the gaps were themselves not evenly distributed within the images, we adopted a randomisation strategy to understand whether the co-localisation values corresponded to enrichment or exclusion of MSCs on gaps, or whether they were consistent with random distributions of cells on an uneven background of gaps.

To assess the distribution of MSCs relative to gaps, we first manually generated binary masks corresponding to the scaffold region for each image. Initially, all [X,Y] coordinate pairs within this mask were set as ‘valid’. Then, each cell object within this mask was allocated a new pair of [X,Y] coordinates randomly drawn from the set of valid coordinate pairs. This new coordinate pair corresponded to the upper-left corner of the bounding box containing the cell object in its new, randomised location. Pixels within this bounding box were now set as ‘invalid’ for the allocation of new coordinates for the next cell, to avoid aberrant overlap between cell positions. This process was repeated iteratively for each cell within the image to generate a randomised MSC label image, and for each original image, 1000 such randomised label images were generated. For each mask i, a co-localisation coefficient ρ’_MG,i_ was computed, yielding a set of 1000 values recapitulating the distribution of co-localisation coefficients consistent with a random distribution of cells on gaps, which could then be compared to the true coefficient ρ_MG_.

### Statistical analysis

Statistical analysis of the data was performed using Prism Version 6 software (GraphPad). Data is presented as the mean ± standard deviation. Statistical significance was assessed using one-way ANOVA test was used to compare the data points. P values used for statistical analysis are indicated in the corresponding figure legends.

## Supplemental figure legends

**Supplemental Figure 1. Optimisation of the scaffold imaging window implantation. (A)** Representative flow cytometry dot plots showing human haematopoietic cells, MSCs and mouse endothelial cells at week 5 after the window implantation. **(B)** Representative characterisation of fluorescence detected in transduced MSC and CD34^+^ haematopoietic stem and progenitor cells used in this study.

**Supplemental Figure 2.** Schematic of optical configuration and light path of the two-photon microscope used in this study. Solid laser lines denote excitation whereas the broken lines denote emitted fluorescence light.

## Supplemental video legends

**Supplemental video 1.** Mouse demonstrating normal ambulatory behaviour immediately following surgical implantation. The animal shows coordinated movement and exploratory activity, indicating good post-operative recovery and tolerance of the procedure.

**Supplemental video 2.** 3-Dimensional reconstruction of humanised scaffold with human MSCs (cyan), haematopoietic cells (red) and collagen (blue). Mice were injected with Texas red labelled 70kDa dextran (yellow) immediately prior to imaging.

**Supplemental video 3.** Time-lapse imaging of a single field of view within the humanised niche, showing dynamic interactions between human haematopoietic cells (Red), MSCs (Turquoise), and the vascular network (Yellow). A fast-moving haematopoietic cell (Red) can be seen actively migrating away from the vessel structures over time.

**Supplemental video 4.** Time-lapse imaging of a single field of view within the humanised niche, showing dynamic interactions between human haematopoietic cells (red), MSCs (turquoise), and the vascular network (yellow). A haematopoietic cell (red) is observed engaging with the vessel structures for an extended period, highlighting sustained cell–vessel interactions within the niche.

## Conflict of interest statement

The authors have declared that no conflict of interest exists.

## Author Contributions

Colin D.H. Ratcliffe and Syed Mian: Performed experiments, analysed data, and wrote the manuscript.

Giovanni Giangreco and Khadidja Habel: Performed experiments.

Alix Le Marois: Analysed data.

Erik Sahai and Dominique Bonnet: Supervised the research, analysed data and reviewed the manuscript.

All authors approved the manuscript.

## Acknowledgments

We would like to thank Kristoffer Riecken for providing us the LeGO plasmids. We would also like to thank staff in the science technology platforms at the Francis Crick Institute including, Cell Sciences, Flow Cytometry, Advanced Light Microscopy, Biological Research Facility and the Making lab. Specifically, we would like to thank Oscar Zabaco Natali, Christina Dix and Donald Bell. This work was supported by the Francis Crick Institute which receives its core funding from Cancer Research UK (CC2027, CC2040), the UK Medical Research Council (CC2027, CC2040), the Wellcome Trust (CC2027, CC2040) to DB and ES, respectively. **C.D.H.R.** is a recipient of a Bourse postdoctorale from the Fonds de recherche du Québec –Santé (https://doi.org/10.69777/273104), as well as an EACR-AstraZeneca Postdoctoral Fellowship. S.A.M is the recipient of MDS foundation young investigator award.

## Disclosures

E.S. reports grants from Novartis, Merck Sharp Dohme, AstraZeneca and personal fees from Phenomic outside the submitted work.

