## Supplementary figures and images for "Intravital imaging uncovers remodelling of humanised bone marrow-like niches"

### Supplemental Figure 1

#### Supplemental Figure 1.

A

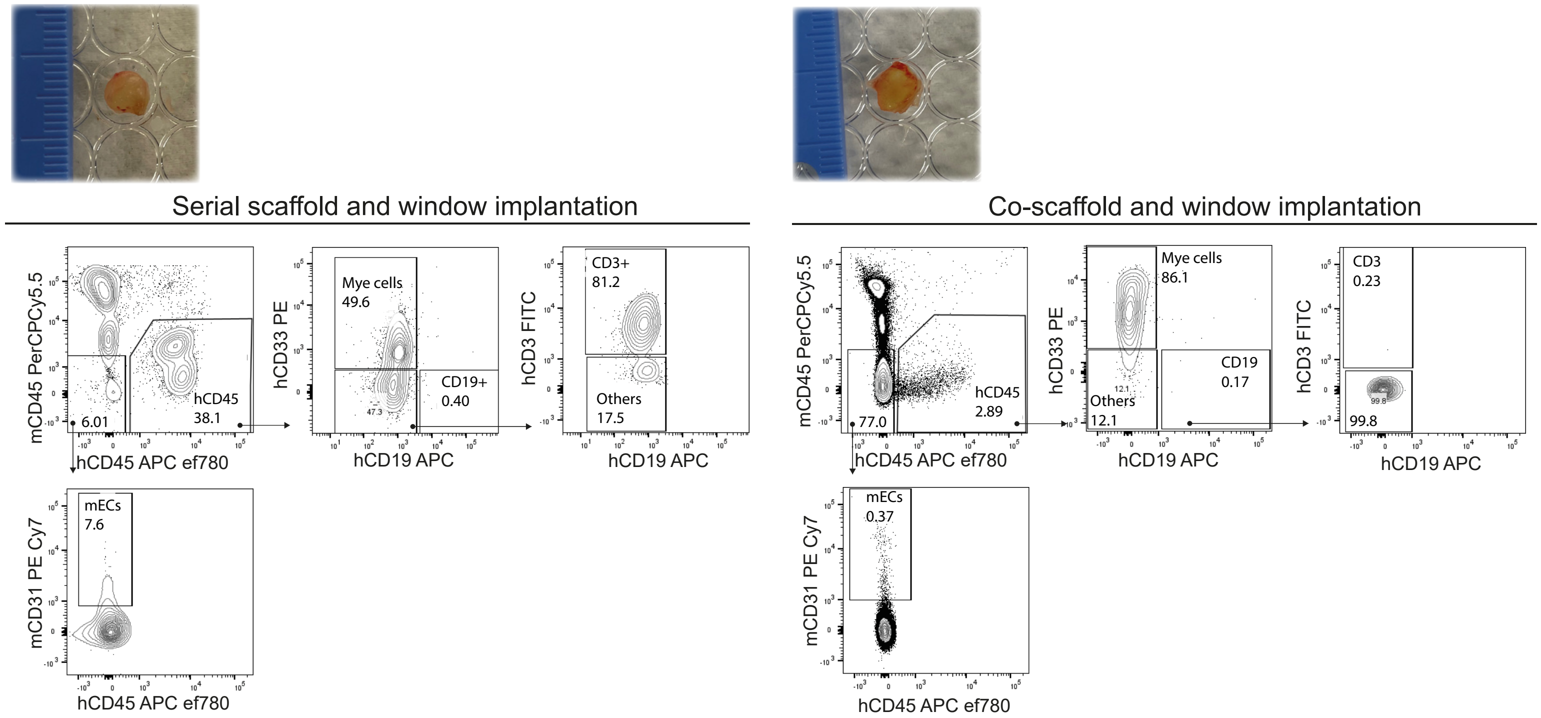


B

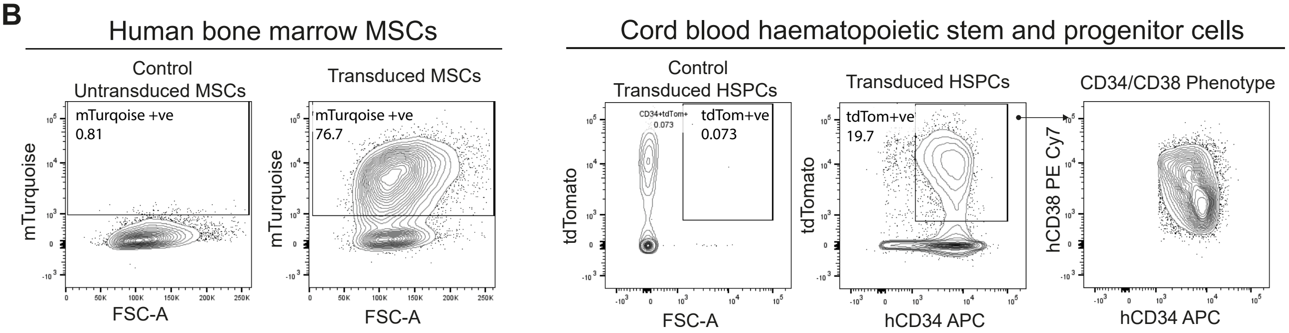

### Supplemental Figure 2

**Supplemental Figure 2**

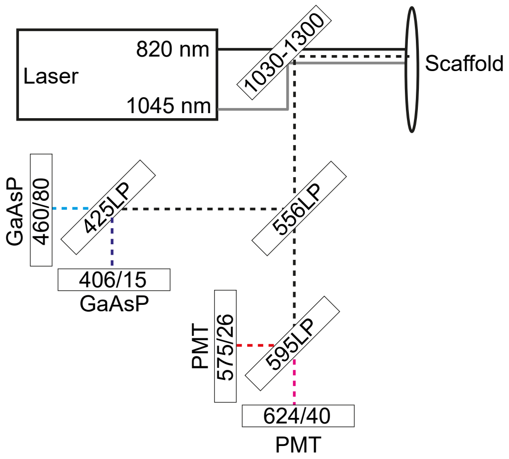
